# Comparative Analysis of Lysine-Specific Peptidases for Optimizing Proteomics Workflows

**DOI:** 10.1101/2024.10.18.619105

**Authors:** Leander van der Hoeven, Maico Lechner, Cristina Hernandez-Rollan, Tanveer S. Batth, Jesper V. Olsen

## Abstract

This study presents a comparative analysis of three LysC homologues from Achromobacter lyticus, Pseudomonas aeruginosa, and Lysobacter enzymogenes for mass spectrometry-based proteomics. Utilizing a protein aggregation capture (PAC) workflow with HeLa cell lysate, we assessed the enzymes’ cleavage specificity, digestion efficiency, and performance across various experimental conditions. Results showed that while all three homologues exhbihited high cleavage specificity at lysine residues, A. lyticus LysC outperformed the two others with its superior peptide identification, digestion efficiency, and protein coverage, especially at short digestion times. Combination of A. lyticus LysC and trypsin demonstrated that importance of LysC for signifancanlty minimizing missed cleavage rates in tryptic digests. This study underscores A. lyticus LysC’s potential as an optimal choice for enhancing mass spectrometry-based proteomics.

## Introduction

Mass spectrometry (MS) is a pivotal analytical technique extensively utilized in proteomics research for the identification and quantification of proteins and their modifications. It is employed in both targeted and exploratory proteomics to analyze proteins and peptides from various biological sources, such as cells, tissues, and fluids like blood or urine.^1^ This technology is also increasingly applied in fields like drug development, food safety, and environmental analysis.^2^ The core strength of MS lies in its ability to provide precise, sensitive, and reproducible measurements of biomolecular compositions.^3^ A typical proteomics workflow involves several stages, including protein digestion into peptides, which are amenable to liquid chromatography-tandem mass spectrometric (LC-MS/MS) analysis.^4^ The complexity of these workflows necessitates meticulous handling and use of highly sequence-specific enzymes to minimize technical variability to ensure reproducibility and quantitative results. Lysine-specific peptidases (LysC) are essential tools in proteomics workflows, enabling efficient digestion of proteins into smaller peptides for MS-based identification and quantification.^5^ LysC is a serine protease that specifically cleaves peptide bonds at the carboxyl side of lysine residues and is commonly used in combination with Trypsin, a peptidase that cleaves C-terminal to arginines and lysines. Unlike Trypsin, LysC is more resistant to denaturants such as urea^6^ and guanidine^7^ and can also cleave after proline. LysC has therefore often been used to predigest protein mixtures under denaturing conditions, whereby proteins are unfolded, allowing Lys-C to access and cleave lysine residues more effectively. This step is crucial for ensuring a more comprehensive breakdown of protein structures and digesting proteolytically resistant proteins effectively before diluting out the denaturants prior to the addition of Trypsin. This combination of peptidases is used to enhance digestion efficiency and reduce missed cleavages at lysine residues.^8^ While LysC-Trypsin combination is currently the gold standard for most MS-based workflows, the performance of different LysC sequences has not been directly compared. Additionally, the properties and efficiency of LysC under different conditions have not been evaluated. LysC was initially discovered in 1970 in the bacterium *Lysobacter enzymogenes* but was soon identified in other bacterial species, including *Achromobacter lyticus* and *Pseudomonas aeruginosa*.^9,10,11^ Efficient protein digestion by proteases such as LysC is influenced by several factors, including the choice of enzyme, sample preparation, and digestion conditions such as pH, temperature, and choice of buffers, among others. However, the specific digestion properties such as the efficiency of different LysC protein sequences under varying conditions have not been directly compared. Therefore, a deep dive could not only improve robustness, but also help improve conditions for a best practice in peptide quantitation. While these sequences share some similarities in amino acid composition, especially *L. enzymogenes* and *A. lyticus*, as we demonstrate, they differ in some of their digestion properties. The structural differences among the LysC sequences are crucial in defining their biochemical properties and interactions with substrates and inhibitors, ultimately impacting their functionality and application in various biotechnological contexts. However, to date, there has not been a comprehensive study comparing these sequences and their impact on protein digestion in MS-based experiments. In summary, this study highlights the importance of optimizing LysC sequences and sample digestion for improving proteomic workflows. *A. lyticus* LysC demonstrates superior performance in terms of total peptide identifications and cleavage efficiency. The optimal combination of LysC and Trypsin is recommended for optimal proteomic analysis.

## Material & Methods

### Materials

LysC sequencing-grade recombinant enzymes from *Achromobacter lyticus* (KPL0033), *Lysobacter enzymogenes (KPL0013)*, and *Pseudomonas aeruginosa (KPL0022)* were kindly provided from KPL ApS, Copenhagen, Denmark.

### Cell culture

Immortalized human epithelial cervix carcinoma adherent cells (HeLa), were grown in 500 cm^2^ dishes in DMEM (Gibco, Waltham, MA, USA) media containing 2 mM L-glutamine supplemented with 10% fetal bovine serum (Gibco, Waltham, MA, USA), 100 U/mL penicillin (Life Technologies, Carlsbad, CA, USA), and 100 μg/mL streptomycin in a 37 °C incubator supplemented with 5% CO2.

### Sample preparation

Cells were lysed with 1 mL of boiling lysis buffer (4% SDS; 100 mM Tris pH 8.5, 5 mM TCEP, and 10 mM CAA) on plate and scraped, followed by incubation at 95 °C, for 10 min with mixing (1000 rpm). Lysates were sonicated with a tip probe (3 minutes, 3 seconds on, 2 seconds off, 100% amplitude, Fisherbrand™ probes Model 120 Sonic Dismembrator). Protein concentration was calculated using the BCA assay and tryptophan assay.^12,13^ For all experiments, 10ug of protein Lysate was digested per sample. Digestion was done following the Protein Aggregation Capture protocol as previously described Batth et al.^14^ Briefly, proteins were aggregated on magnetic microbeads (MagResyn Hydroxyl beads) by adding acetonitrile (ACN) to a final concentration of 70%. Beads were washed once with 100% ACN and once with 70% ethanol. 50 mM HEPES digestion buffer at pH 8.5 was added to all samples prior to the addition of proteases. Unless stated otherwise, a typical protease-to-protein ratio of 1:100 was employed for all the digestion. The digestion reaction was quenched by adding trifluoroacetic acid (TFA) to a final concentration of 1%. Thereafter, samples were loaded into Evotips for subsequent MS analysis.^15^

### LC-MS/MS analysis

Digested peptides were separated on a EV-1109 column (PepSep, 8 cm × 150 µm, beads 1.5 um) and EV-1087 emitter (fused silica, 20 µm) with the Evosep One LC system. The column was interfaced online using an EASY-Spray™ source with the Orbitrap Exploris 480 MS (Thermo Fisher Scientific, Bremen, Germany) using Xcalibur (tune version 4.0 or higher). In all samples, spray voltage was set to 1.8 kV, funnel RF level at 40, and heated capillary temperature at 275 °C. All experiments were acquired using a data-dependent acquisition (DDA) top 20 method with a 60 samples per day (SPD) LC gradient.

### Data analysis

All Thermo raw files were searched with the MaxQuant software suite 2.4.13.0 at default settings or LysC/P semi-specific search using a human database (Uniprot reference proteome 2023 release, 20,598 entries) in addition to a contaminants FASTA file containing porcine-Trypsin and the LysC sequences utilized in this study.

The output files were analyzed using Rstudio with the following packages, dplyr (1.1.2), readr (2.1.5), tidyr (1.3.1), ggplot2 (3.5.1), ggrepel (0.9.3), stringr (0.9.3), ggfortify (0.4.17), ggpattern (1.1.1), biomaRt (2.56.1), httr (1.4.7), ggseqlogo (0.2) and jsonlite (1.8.8).

## Results

To identify the best LysC for proteomics research, we analyzed the cleavage specificity and digestion efficiency of three LysC homologues from *Lysobacter enzymogenes, Achromobacter lyticus*, and *Pseudomonas aeruginosa*, respectively. These three LysC homologues and variations of them are the main LysC enzymes used in proteomics. A protein sequence alignment of the LysC enzymes from *A. lyticus, P. aeruginosa*, and *L. enzymogenes* reveals conserved regions essential for enzymatic function, especially the triad essential for activity consisting of residues of Ser194, His57, and Asp113 of *A. lyticus* and *L. enzymogenes* respectively, and His72, Asp122, and Ser196 for *P. aeruginosa*, and variations that may influence specificity and efficiency (Figure 1A). To benchmark the performance of these three enzymes against each other, we applied them in an experimental sample preparation workflow based on PAC of HeLa cell lysates and subsequent on-bead digestion ranging from 30 to 960 minutes and analyzed the resulting peptide mixtures by online LC-MS/MS (Figure 1B). Each digestion condition was analyzed in four workflow replicates to assess reproducibility.

**Figure 1.**
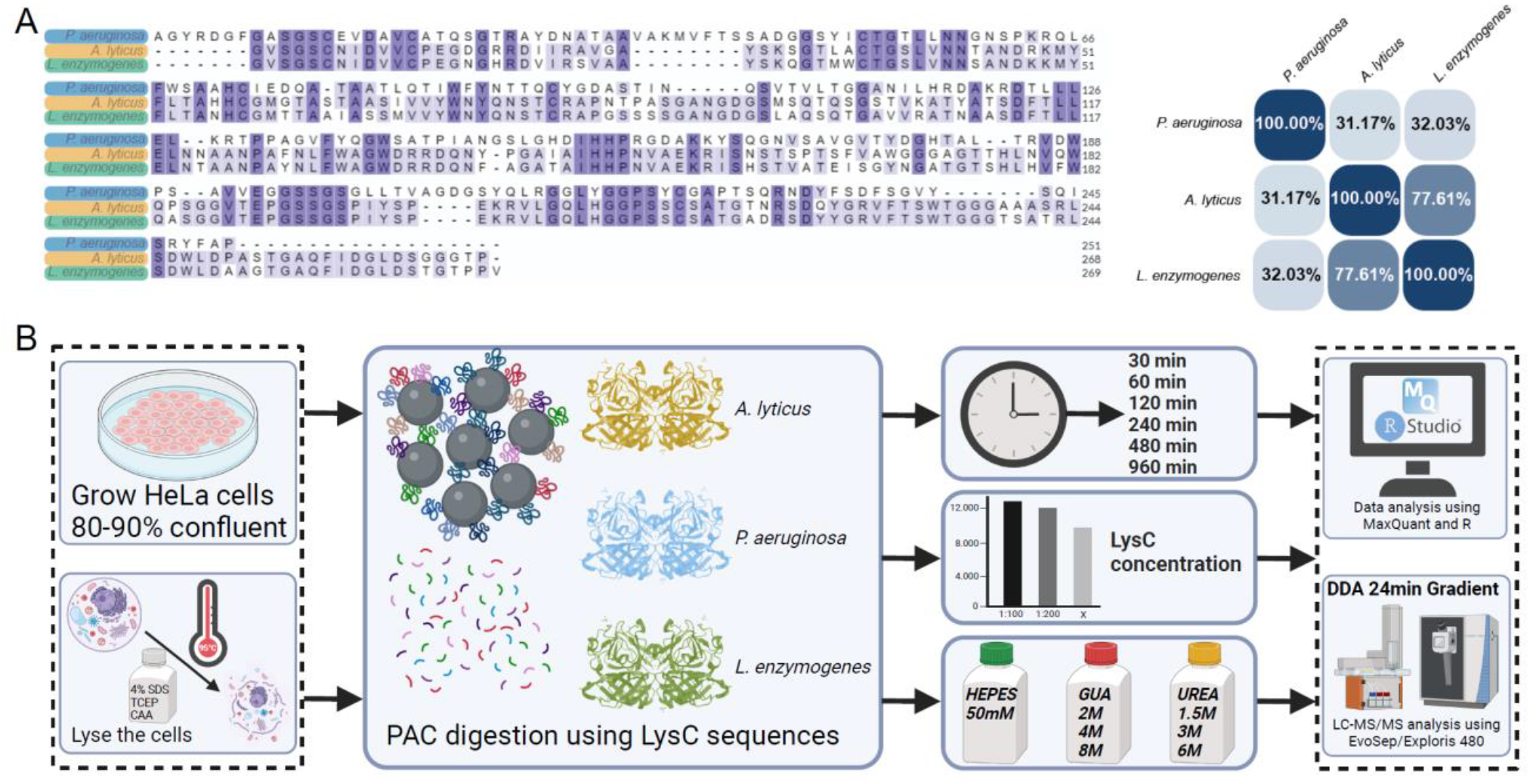
Sequence analysis of the three LysC enzymes used and workflow representation of the study. A) Left panel: sequence alignment of the catytic regions of LysC enzymes from P. aeruginosa, A. lyticus, and L. enzymogenes. Conserved regions are highlighted in blue. Right panel: Homology score in percentage between the three LysC enzymes. B) Schematic representation of the workflow used to compare the three enzymes in this study.

First, we evaluated the efficiency of the three LysC homologues by analyzing the total number of HeLa peptides plotted as a function of digestion time. This revealed that LysC from *A. lyticus* consistently identified more peptides, especially in the early time points (Figure 2A, left panel), indicating superior performance in complex samples, especially when compared to *L. enzymogenes*. This was also the case when analyzing the summed total peptide intensities as a proxy for protease efficiency (Figure 2A, middle panel). The trend was largely the same when evaluating the three LysC homologues based on the total HeLa protein numbers identified as a function of digestion time with the exception that with the overnight digestion they all identified an almost comparable number of proteins (Figure 2A, right panel). To assess if the observed difference in protease efficiency is linked to their cleavage activity, we analyzed this as a function of digestion time. Cleavage efficiency represented by the percentage of peptides with missed lysine cleavage sites confirmed that *Achromobacter lyticus* exhibits the highest enzymatic efficiency with the lowest level of missed cleavages observed, reaching close to 90% efficiency at early time points and 92% of detected peptides with zero missed cleavages after an overnight digestion (Figure 2B, left panel). Conversely, LysC homologues from both *Pseudomonas* and *Lysobacter* achieved aabout 70% cleavage efficiency at best, respectively (Figure 2B, middle and right panels).

**Figure 2.**
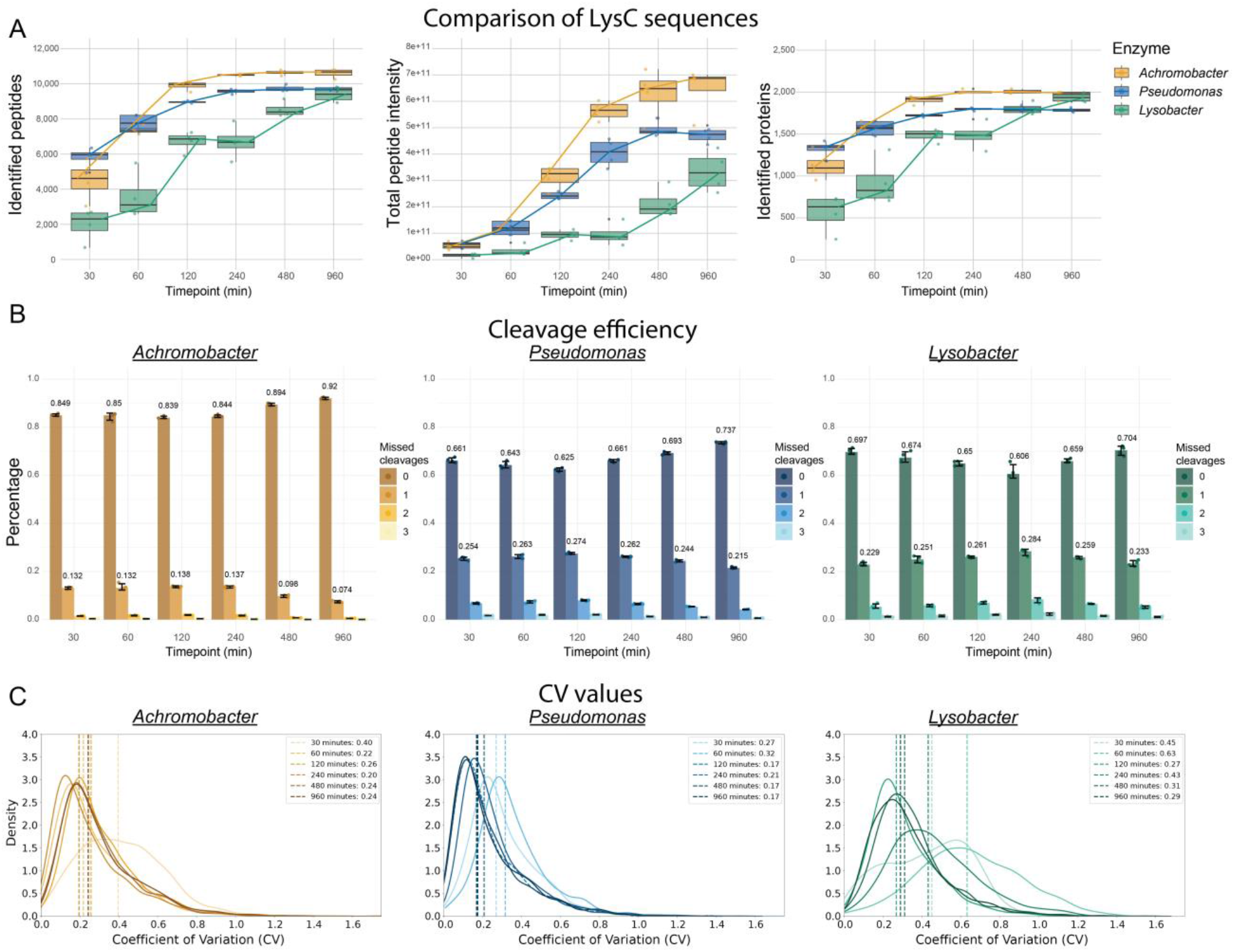
Digestion efficiency evaluation for the three LysC homologues. A) Comparison of the identified peptides, total peptide intensity, and number of identified proteins over a digestion time course ranging from 30-960 minutes for the three LysC homologues. B) Cleavage efficiency of the three LysC sequences is measured by the percentage of missed cleavages. C) Coefficient of variation (CV) values across different digestion time points.

Finally, to evaluate the quantitative performance of the three LysC homologues, we analyzed the coefficient of variation (CV) values for the identified peptides across the three replicates. This analysis showed that *A. lyticus* and *P. aeruginosa* display similar CV values, while *L. enzymogenes* shows higher and more variable CVs, indicating more consistent performance over time for both *A. lyticus and P. aeruginosa* (Figure 2C).

To evaluate the cleavage specificity of the three LysC homologues, we reanalyzed the overnight digestion files allowing for semi-specific peptide sequence matches. Cleavage specificity analysis of all identified peptide sequences showed that all three LysC homologues are highly specific, withmore than97% of all cleavages at the C-terminal of lysine residues and no apparent secondary preferences (Figure 3A). To identify any bias for the LysC homologues against specific amino acids in the proximity of lysine residues in substrate proteins, we performed a sequence motif enrichment analysis of the amino acid sequences surrounding all missed cleaved lysine sites identified. Such a sequence motif analysis can highlight sequence-specific patterns contributing to missed cleavages, suggesting structural features affecting enzyme-substrate interactions. The analysis revealed an enriched sequence pattern for missed cleavage lysine sites with bias against glutamic acid (E) and additional lysine (K) in close proximity of cleavage sites (Figure 3B). This enriched motif was highly similar for all three LysC homologues.

**Figure 3.**
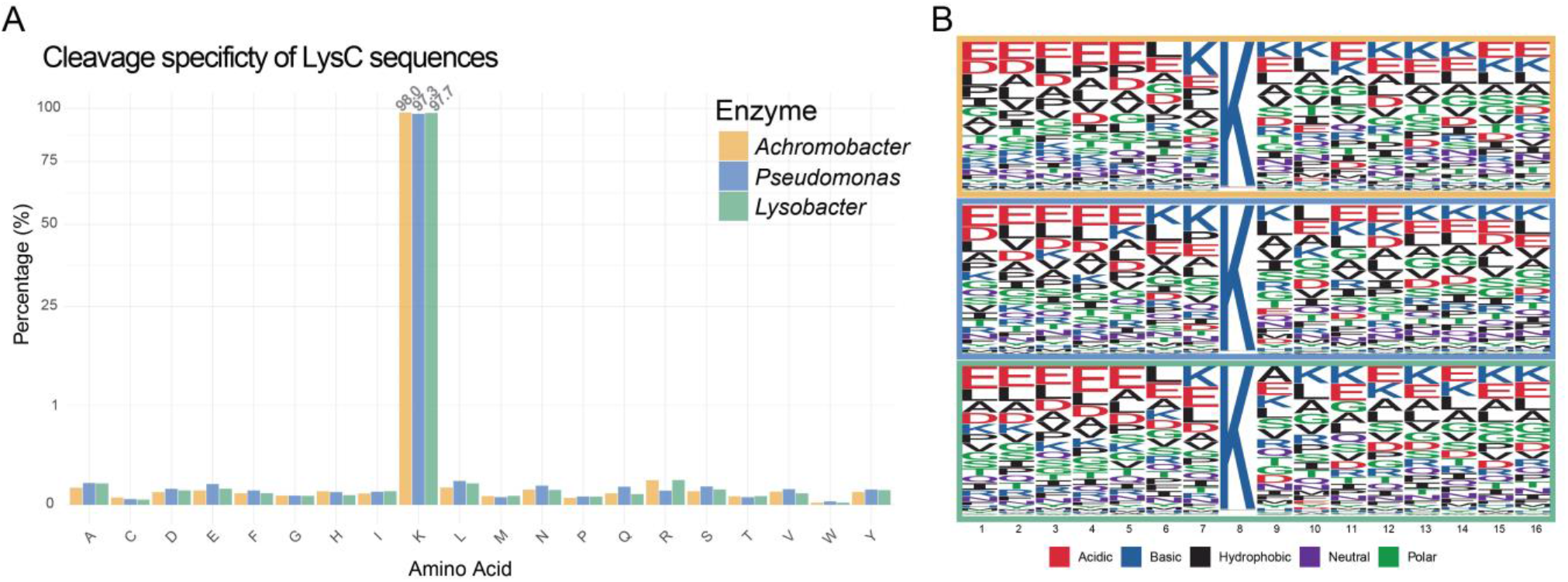
Catalytic specificity evaluation for the three LysC homologues. A) Percentage evaluation of the amino acid cleavage specificity analysis of LysC enzymes. B) Motif analysis of peptides containing missed cleavages.

To compare the HeLa proteins identified by the three LysC homologoues in more detail, we first looked at the overlap between them. A Venn diagram displayed the overlap and unique proteins identified by each LysC sequence (Figure 4A). *A. lyticus* identifies the most unique proteins, underscoring its enhanced coverage. To analyse their digestion patterns closer. we examined the sequence coverage achieved by the three LyCs of a highly abundant protein, PGK1 (Figure 4B). This comparison showed that all three enzymes accomplished high sequence sequence but *A. lyticus* achieved the highest with fully cleaved peptides compared to its two counterparts, where a large proportion of the peptides identified showed at least one missed cleavage.

**Figure 4.**
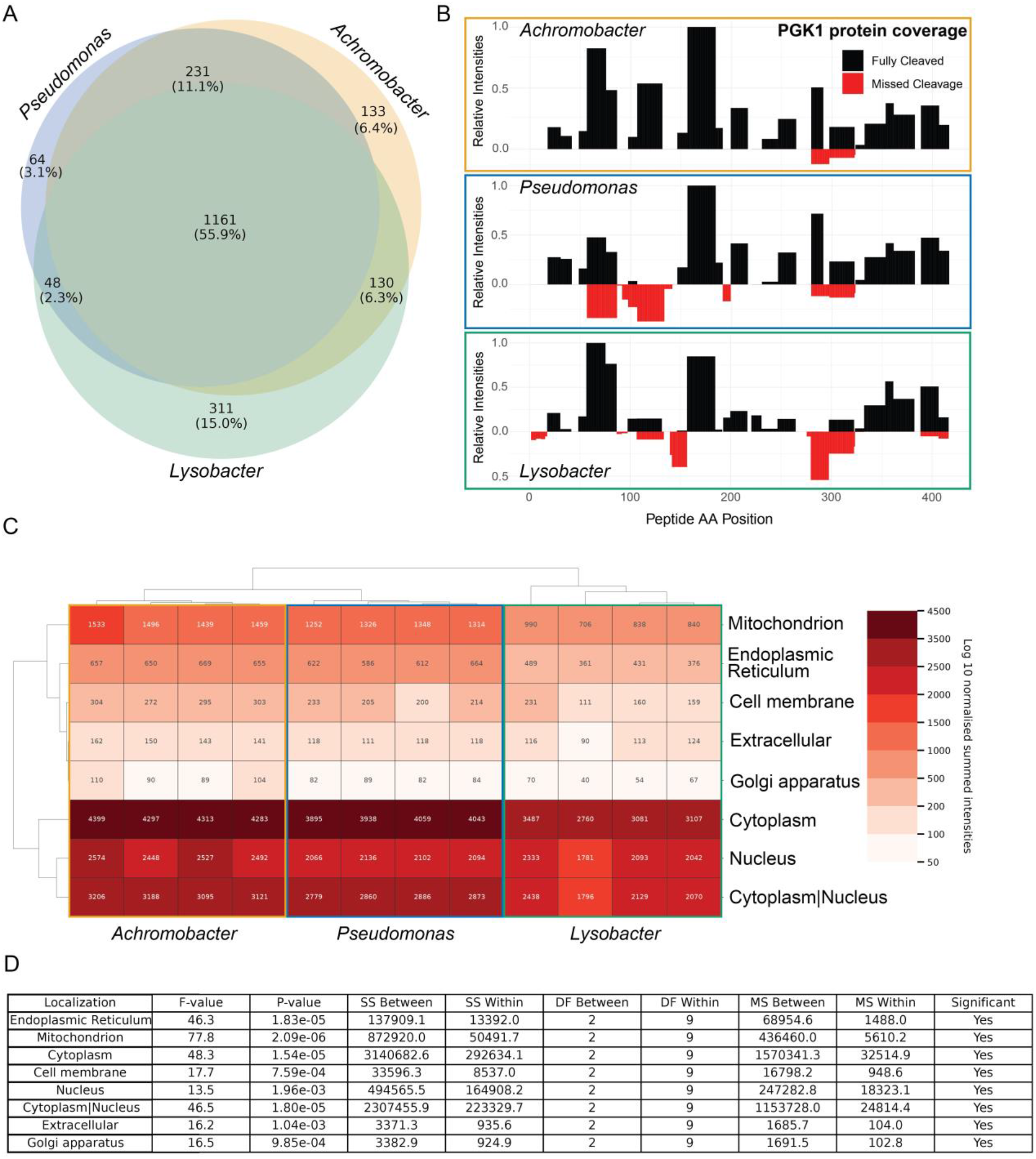
Comparison of protein identification, digestion patterns, and subcellular localization across different LysC enzymes. A) Venn diagram of proteins identified by each LysC enzyme, showing the overlap and the unique proteins shared between them. B) Protein coverage analysis of the PGK1, comparing the digestion patterns of the three LysC sequences. The relative intensities of the identified peptides are plotted based on their positions within the protein. Additionally, the fully cleaved peptides are shown in relation to those showing a missed cleaved. C) Heatmap of protein subcellular localization for each LysC enzyme, revealing from which compartment they originate. D) Heatmap depicting the log10 normalized protein intensities connected to several localized cell substructures. Both cellular localization and LysC types were clustered using a hierarchical clustering algorithm based on Euclidean distance similarity.

To assess potential biases in protein classes identified by the three LysC homologues, we grouped the identified proteins by their subcellular localization based on data from the DeepLoc 2.0 project.^15^ We performed unsupervised hierarchical clustering of log10-transformed intensities of all identified proteins by their cellular compartment across the four replicates for each LysC enzyme and visualized it in a heatmap (Figure 4C). The heatmap exhibited distinct patterns of protein abundance across different subcellular components. The hierarchical clustering revealed that the replicates of each LysC type group closely together, indicating high consistency within each LysC type. The clustering on the subcellular localisation axis showed two subgroups emphasizing a difference between proteins localized in the nucleus and cytoplasm. The second cluster includes the subcellular components of the mitochondrion, endoplasmic reticulum, cell membrane, and extracellular proteins. Intensities indicated that LysC from *Achromobacter lyticus* showed most similarities with *Pseudomonas aeruginosa. Lysobacter enzymogenes* showed overall lower protein intensities for all localisations compared to the two other LysC enzymes, which was coherent with the overall reduced performance of *Lysobacter enzymogenes* displayed in Figure 2A. To further evaluate if there is statistical significance between the 3 LysC types, we performed a one-way ANOVA statistical test for each subcellular localisation group comparing the sum of squares (SS), mean squares (MS) and degrees of freedom (df) between and within each group (Figure 4D). Based on this, input F-values was calculated and compared to the tabulated critical F-values. The results are presented as p-values indicating a statistically significant difference between the variance of a localisation if the p-values is below a significance threshold of 0.05. The F-values and corresponding p-values indicated significant differences across all localizations, with the highest F-value observed in the mitochondrion (77.8, p = 2.09e-06), suggesting a substantial variation of the intensities between the three LysC types. The endoplasmic reticulum and cytoplasm also showed high F-values (46.3 and 48.3,respectively), indicating significant differences. The cell membrane and nucleus exhibited lower but still significant F-values (17.7 and 13.5, respectively). The sum of squares between groups is notably high in the cytoplasm and nucleus, reflecting substantial variability in these regions. The mean squares between groups are consistently higher than within groups, reinforcing the significant impact of the different LysC types and suggesting a good reproducibility between technical replicates. All of the above experiments described in figure 1-4 consistently demonstrated that A. lycitus as the superior LysC compared to the two others.

As the most frequent use of LysC in proteomics is in combination with trypsin, we next tested the impact combining the A. lyticus LysC with sequencing grade modified trypsin for overnight tryptic digestion using a fixed trypsin-to-protein ratio of 1:50. LysC in different protease-to-protein ratios was tested either as a predigestion step prior to trypsin addition or as a combo with trypsin adding LysC and trypsin at the same time. The impact of LysC addition was evaluated based on the fraction of missed cleavage rates at arginines and lysines and compared to trypsin only digestion (Figure 5A). A LysC predigestion step showed that the optimal protease combination for LysC is 1:100 together with the trypsin-to-protein ratio of 1:50 (Figure 5A, left panel). Predigestion of the protein mixture did not seem to have an effect in reducing the number of missed cleaved peptides compared to adding LysC and trypsin at the same time with all tested combinations showing comparable ∼85% identification of tryptic peptides with zero missed cleavages (Figure 5A, left panel). Conversely, using only trypsin without LysC significantly increases the number of missed cleavages with only ∼70% identification of tryptic peptides with zero missed cleavages. This is especially apparent in the lysine-containing peptides where ∼30% of all identified peptides contain a missed cleavage when using trypsin without LysC, whereas less of difference is observed for arginines (Figure 5A, right panel). This analysis reveals that peptides containing lysines are more prone to missed cleavages during tryptic digests, suggesting that trypsin is suboptimal for cleaving lysines. This demonstrates the importantance of using LysC in combination with trypsin for minimizing missed-cleavage sites in tryptic digests. While there does not appear to be a clear motif suggesting a cause for the missed cleavages, there is a larger concentration of glutamic and aspartic acids, leucines, lysines and prolines near the lysine sites that are cleaved by the LysC and trypsin combo but not by trypsin alone (Figure 5B).

**Figure 5.**
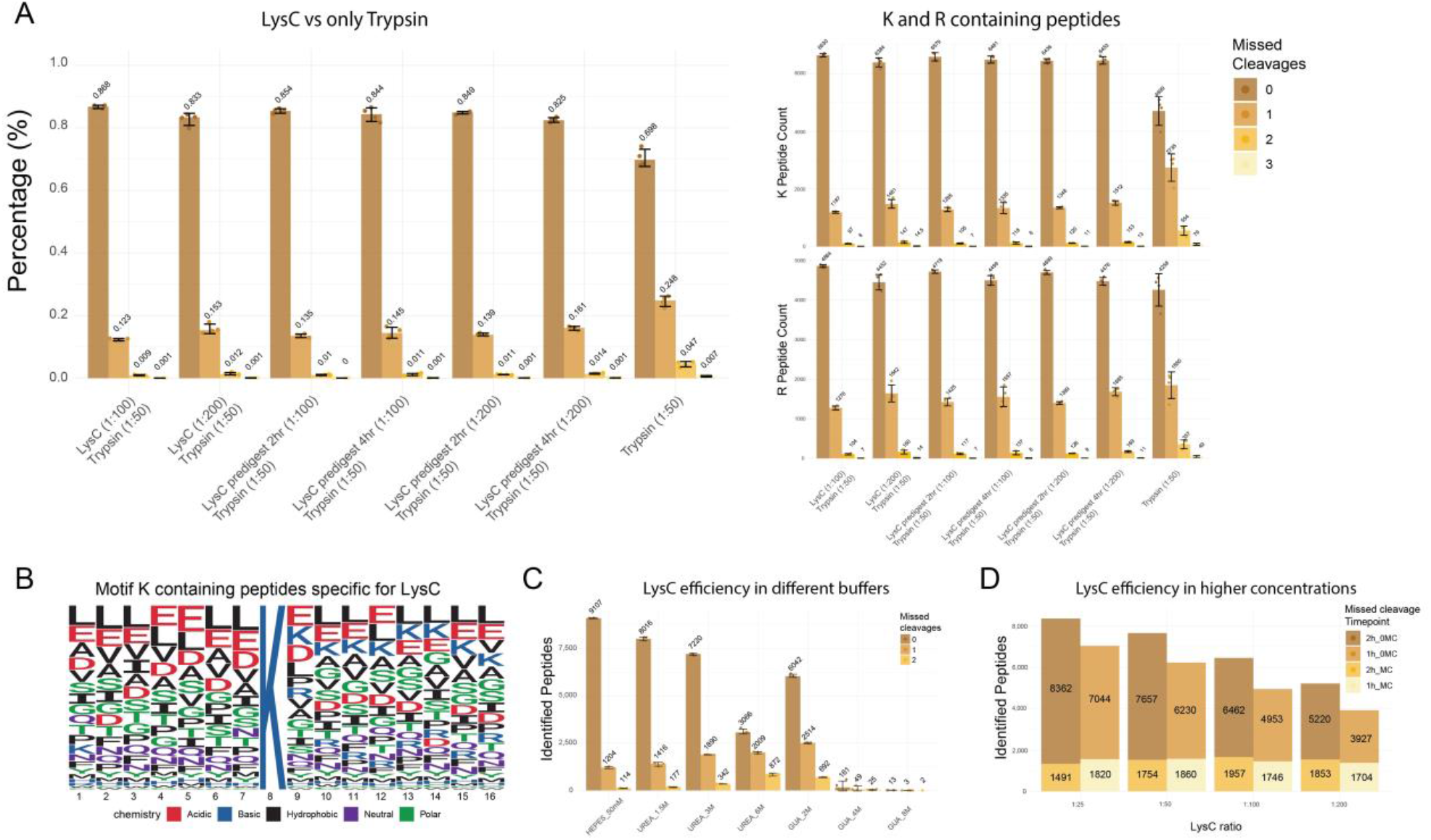
Evaluation of LysC digestion efficiency and cleavage specificity of A. lyticus across varying conditions. A) Comparison of LysC concentrations and digestion methods for the A. lyticus LysC sequences. In the right panel peptides specifically containing Lysine or Arginine are highlighted. B) Motif of K containing peptides containing 0 missed cleavages in LysC/Trypsin combination and containing at least one missed cleavage in Trypsin only. C) Digestion efficiency of A. lyticus under different denaturing buffers. D) Digestion efficiency of A. lyticus at several concentrations.

As LysC is often used to predigest protein mixtures under harsher denaturing conditions than trypsin can tolerate, we next tested the cleavage efficiency of A. lyticus LysC in different denaturing buffers. This analysis showed that digesting a HeLa lysate at varying concentrations of denaturing buffers, the performance of LysC is relatively stable in urea up to 1.5M (Figure 5C). In higher urea concentrations, a similar number of peptides are identified, however, with a markedly increased number of missed cleavage sites. Conversely, LysC is more sensitive to Guanidinium (GUA) with a clear loss in the number of identified peptides, already at the lowest tested concentration of 2M GUA, and with the highest tested concentration leading to close to zero peptides being identified (Figure 5C).

Finally, to identify the best compromise between LysC-to-protein ratio and digestion time, we analysed different combinations of these. When digesting with a higher concentration of *A. lyticus* LysC the digestion time showed an almost linear curve compared to the LysC concentration (Figure 5D). Interestingly, this analysis suggests that a rule of thumb, doubling the amount of LysC leads to the same number of peptides identified in half the digestion time.

In conclusion, while all three LysC homologues effectively facilitate protein digestion in mass spectrometry-based proteomics workflows, *A. lyticus* demonstrates superior cleavage specificity, efficiency, protein coverage, and consistency across the tested conditions. These findings highlight its potential as an optimal choice for enhancing proteomics workflows.

## Discussion

The comprehensive benchmark analysis of three LysC homologues from *A. lyticus, P. aeruginosa* and *L. enzymogenes*, provides valuable insights into their performance in typical mass spectrometry-based proteomics research. This study evaluated the enzymes’ cleavage specificity, digestion efficiency, and overall performance. The results consistently demonstrated that LysC from *A. lyticus* outperformed its counterparts in terms of peptide identification and digestion efficiency, particularly with short digestion times of few hours, suggesting its potential to enhance the sample throughput of proteomic studies, especially when a higher LysC endopeptidase to protein ratio was chosen. This superior performance was further evidenced by its high cleavage efficiency, reaching nearly 90% at early time points and 92% after overnight digestion, with the lowest level of missed cleavages observed among the three LysC sequences. In terms of quantitative performance, *A. lyticus* and *P. aeruginosa* LysC displayed similar and more consistent coefficient of variation (CV) values across replicates compared to *L. enzymogenes*. This consistency is crucial for quantitative proteomics, where reproducibility is paramount. All three LysC homologues exhibited high specificity for lysine residues, with over 97% of cleavages occurring at the C-terminal of lysine. However, a sequence motif analysis revealed a bias against glutamic acid and additional lysine residues near cleavage sites in missed cleavages, which could be valuable for predicting potential missed cleavages and optimizing digestion protocols. The enhanced protein coverage achieved by *A. lyticus* LysC, as demonstrated by the analysis of the PGK1 protein. The subcellular localization analysis revealed distinct patterns among the LysC homologues, with *A. lyticus* showing broader distribution across cellular compartments, particularly in mitochondria and cytoplasm. Investigating the use of LysC in combination with trypsin at different protein-to-protease ratios and predigestion steps, provided valuable insights for optimizing proteomics workflows. Trypsin digestion without LysC yielded significantly higher missed cleavage rates, especially at lysine residues, emphasizing the importance of LysC for enhancing tryptic digestion. The study found that the direct combination of LysC (1:100) and trypsin (1:50) yielded optimal results without the need for a predigestion step, potentially streamlining sample preparation protocols. Additionally, the study demonstrated that *A. lyticus* LysC stability in varying concentrations of denaturing buffers, offering flexibility in protein extraction and digestion buffers, particularly when dealing with challenging protein samples. While this study provides a comprehensive comparison of the three most popular LysC homologues, several avenues for future research emerge. These include structural analysis to understand the molecular basis for varying efficiencies, evaluation of performance across diverse sample types, exploration of potential synergistic effects with other proteases, and application in large-scale, quantitative proteomic studies to assess its impact on proteome coverage in real-world applications.

In conclusion, this study provides compelling evidence for the superior performance of *A. lyticus* LysC in proteomic workflows. Its high efficiency, specificity, and consistency make it an excellent choice for researchers aiming to enhance the depth and quality of their proteomic analyses. The insights gained from this comparative study not only inform the selection of proteases for current research but also pave the way for further optimization and innovation in proteomic methodologies.

## Acknowledgements

We kindly thank KPL ApS, Copenhagen, Denmark.for providing us LysC sequencing-grade recombinant enzymes from Achromobacter lyticus, Lysobacter enzymogenes, and Pseudomonas aeruginosa (KPL0022) were kindly provided from KPL ApS, Copenhagen, Denmark. Work at The Novo Nordisk Foundation Center for Protein Research (CPR) is funded in part by a donation from the Novo Nordisk Foundation (NNF14CC0001 and NNF21OC0072070). J.V.O. is also funded by Novo Nordisk a/s (CELFFI-2022-002843).

